# Modulation of neural variability: Age-related reduction, GABAergic basis, and behavioral implications

**DOI:** 10.1101/2022.09.14.507785

**Authors:** Poortata Lalwani, Thad A. Polk, Douglas D. Garrett

## Abstract

Moment-to-moment neural variability has been shown to scale positively with the complexity of stimulus input. However, the mechanisms underlying the ability to align variability to input complexity are unknown. Using a combination of computational modeling, fMRI, MR spectroscopy, and pharmacological intervention, we investigated the role of aging and GABA in neural variability during visual processing. We found that participants expressed higher variability when viewing more complex stimuli. Such variability modulation was associated with higher baseline visual GABA levels and was reduced in older adults. When pharmacologically increasing GABA activity, we found that participants with lower baseline GABA levels showed higher drug-related increase in variability modulation, consistent with an inverted-U account. Finally, higher baseline GABA and variability modulation were jointly associated with better visual-discrimination performance. These results suggest that GABA plays an important role in how humans utilize neural variability to adapt to the complexity of the visual world.

## INTRODUCTION

Our sensory modalities are constantly processing myriad inputs that differ in complexity and familiarity across various dimensions. And yet, our neural networks can accommodate and process such heterogeneity. It has been postulated that an efficient neural network may do this by modulating neural dynamics to align with the complexity of input stimuli; neural dynamics would be increased to process more complex stimuli, but compressed to process less complex stimuli^1–3^. One neural measure that scales with stimulus complexity and captures this ability is moment-to-moment variability in the blood oxygen level-dependent signal (SD_BOLD_). Recent research on older adults found that the ability to upregulate visuo- cortical SD_BOLD_ when viewing more complex, feature-rich visual stimuli characterized those with better visuo-cognitive performance^3^. These results suggest that upregulation of variability in response to the increased complexity of stimulus input (or 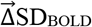) may be an index of the flexibility of an individual’s neural system and predict individual differences in sensory processing. Moreover, previous research suggests that the modulation of variability in response to task demands is reduced in older adults compared to younger adults^4–6^ and that this reduction is associated with poorer behavioral performance^4,6–10^. We thus hypothesized that 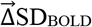 (expected to scale positively with visual stimulus complexity) would also be lower in older adults and that this reduction would be linked with poorer visual discrimination. But what is the basis of such age-based (and individual) differences in 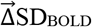 in the first place?

Gamma-aminobutyric acid (GABA), the brain’s major inhibitory neurotransmitter, is one plausible candidate. GABA has been associated with cortical plasticity^11–13^, with pattern complexity^14^, and with the dynamic range^15,16^ of neural networks. Computational work has found that altering the strength of inhibitory connections in artificial neural networks can dramatically affect moment-to-moment dynamics and the number of different states that networks can sample^16^. Similarly, decreasing GABA activity pharmacologically in healthy young rats and monkeys leads to a decrease in network dynamics and reduces the number of states visited by the cortical network^15^. In humans, GABA levels are lower in older adults^17–28^ and in recent work, we found that overall cortical resting-state SD_BOLD_ can be increased in older, poorer performing adults by GABA agonism^29^. However, it remains unknown whether GABA provides a principled basis for how humans align brain signal variability to the complexity of visual input, a crucial step for understanding the mechanisms subserving task-relevant variability modulation in the human brain. Under the assumption that temporal dynamic range is required to distinctly represent external stimuli and respond to stimulus complexity^3,30^, it is plausible that boosting GABA in individuals with lower baseline GABA levels should increase 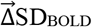, while those with higher baseline levels of GABA would exhibit a decrease in 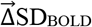, consistent with an inverted-U effect^31^.

Using data from 58 younger (ages 18-25) and 77 older (ages 65-85) adults (see Figure 1 for complete study design), we addressed these various questions using a combination of computational modeling, functional magnetic resonance imaging (fMRI), magnetic resonance spectroscopy (MRS), pharmacological intervention, and behavioral analyses. Specifically, we first utilized a computational model of the visual system (HMAX) to estimate the complexity of visual stimuli seen by participants during fMRI. To examine the GABAergic basis of such variability modulation, we then (1) measured baseline visuo-cortical GABA levels using MRS, and (2) used lorazepam (a GABA_A_ agonist) to causally increase GABA activity, while assessing visual 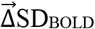 on and off drug. In a different subset of participants, we computed a latent visual discrimination behavioral score across four offline tasks. We tested whether: (1) neural variability scaled in response to stimulus complexity; (2) 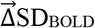 was more modest in older adults; (3) lower 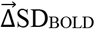 was associated with lower visual GABA levels; (4) pharmacologically increasing GABA activity increased 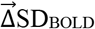 in those with low GABA levels, and; (5) higher GABA and greater 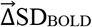 were associated with better visuo- behavioral performance.

**Figure 1:**
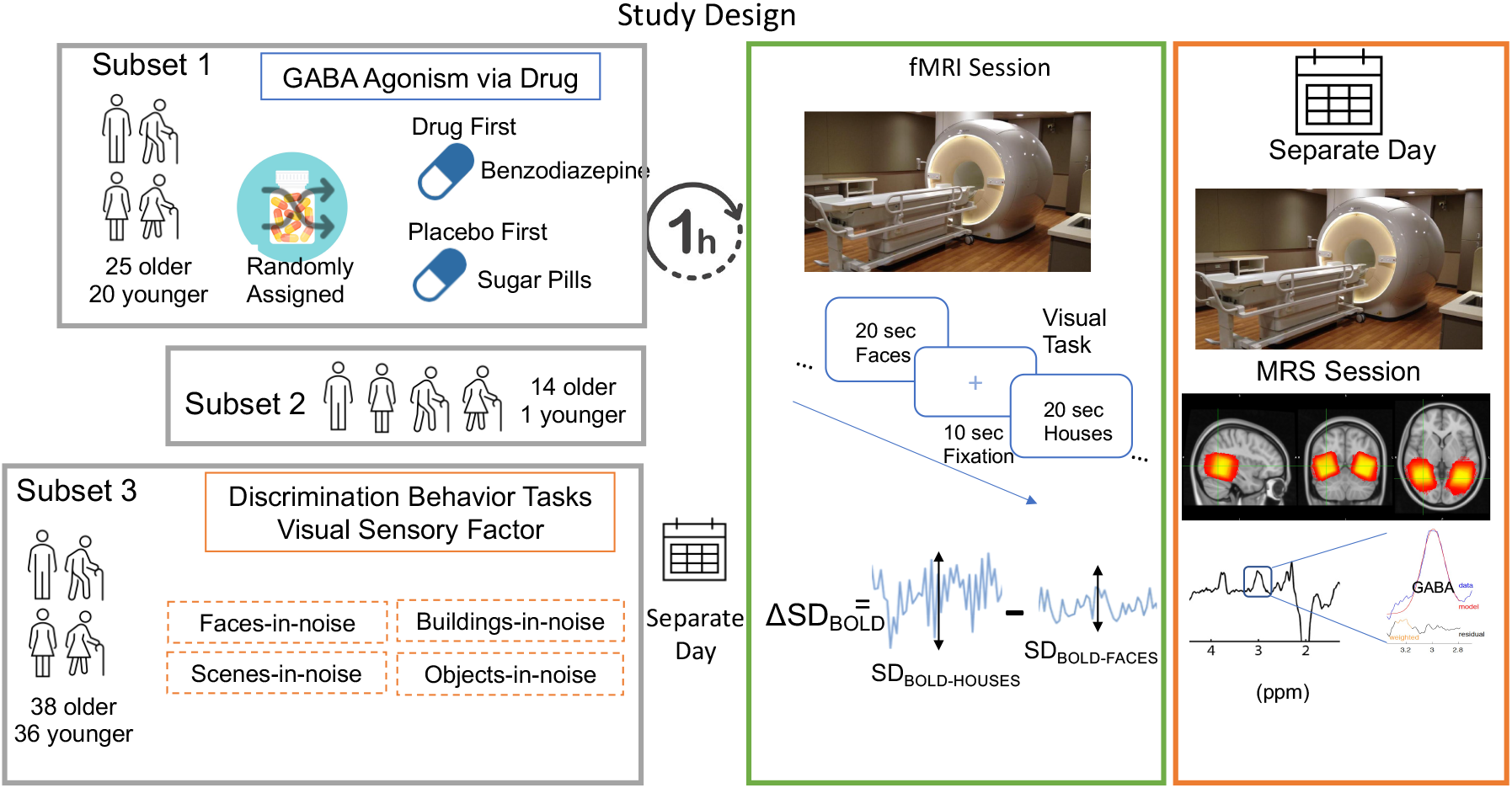
Session Design and Participant Distribution. All participants underwent an fMRI and MRS scanning session on separate days. One subset of participants (25 older and 20 younger) received an additional on-drug fMRI scan on a separate day. The order of on-drug and off-drug fMRI sessions was randomized. A different subset (3) of participants (38 older and 36 younger) completed four visual discriminatory tasks on a separate day before fMRI testing. During the fMRI session participants completed a 6-minute visual task with pseudorandomized 20-second blocks of passively viewed houses and faces interleaved with 10-second fixation blocks as shown in the green panel. Change in variability 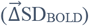 is computed at every voxel as the difference between SD_BOLD-HOUSES_ and SD_BOLD-FACES_. The orange panel shows the MRS voxel overlap across participants with brighter (yellow) indicating maximum overlap and red showing the least, an example spectrum obtained, and that raw GABA+/water is estimated by fitting a Gaussian model to compute the area under the curve of the 3ppm GABA peak.

## RESULTS

### Visual stimulus complexity and variability modulation in younger and older adults

We first estimated the complexity (or “feature-richness”) of our set of visual stimulus categories (faces and houses) by submitting them to a biologically inspired computational model of visual processing (HMAX)^32–35^. The early layers in this model correspond to neurons in primary visual cortex (V1) and the later layers correspond to neurons in extrastriate visual areas (V2/V4). As argued by Garrett et al. (2020), faces may be encoded with a smaller number of perceptual dimensions yet expertly discriminated by humans, while houses are more differentiated stimuli, less constrained in their form. We thus presumed that HMAX would detect houses as a more feature-rich stimulus category than faces, with houses also presumably requiring greater dynamic range (SD_BOLD_) during processing. Indeed, HMAX revealed that houses were more feature-rich than faces in both early (corresponding to V1) and later model layers (corresponding to V2/V4; all t-values > 10). See Supplementary Materials for full model description and results.

To examine whether brain signal variability is modulated in line with visual complexity, we computed the standard deviation of the BOLD signal (SD_BOLD_) during each task condition (houses and faces) within each voxel and participant after extensively pre-processing the fMRI data (see Methods). Using Partial Least Squares (PLS), we found a single significant latent variable (p<0.001) revealing higher SD_BOLD_ when viewing houses than faces (SD_BOLD-HOUSES_ > SD_BOLD-FACES_), particularly in ventral visual cortex (See Figure 2 for spatial extent and Table S1 for cluster details). This task-condition effect on SD_BOLD_ was significant in both younger (t(57) = 3.43, p <0.001, Cohen’s d = 0.45) and older adults (t(76) = 4.15, p <0.001, Cohen’s d = 0.47).

**Figure 2:**
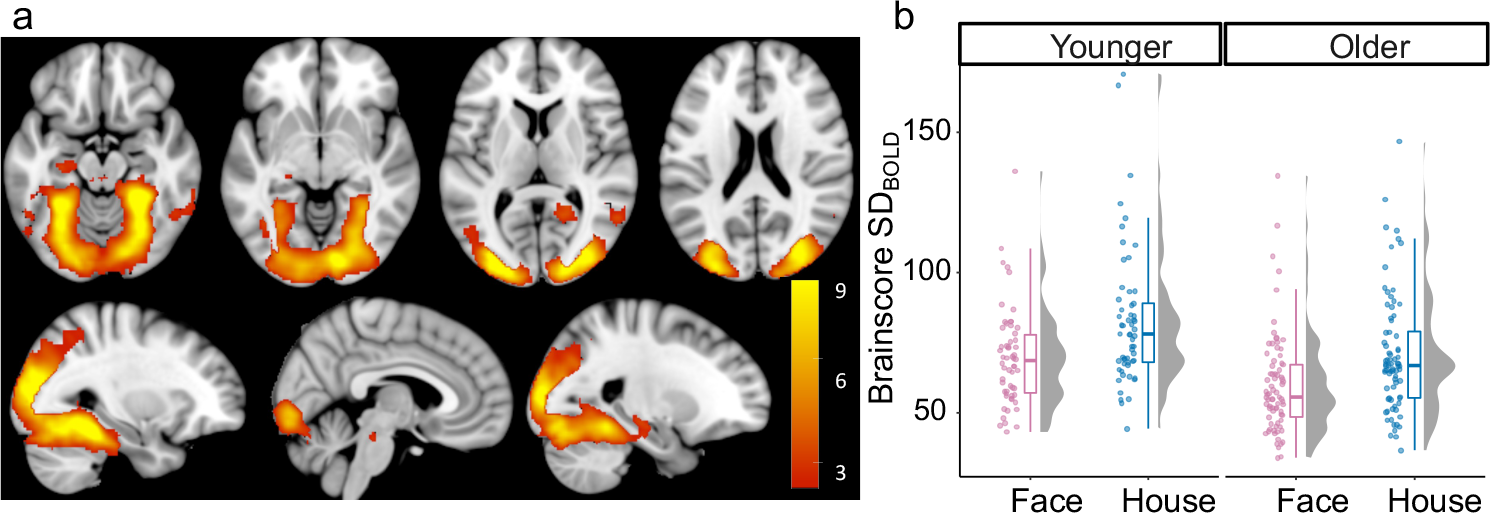
Effect of task-condition on SD_BOLD_. Primary and extended visual cortex show reliably higher variability (Cohen’s f = 0.46) during the house condition (in blue) than face condition (in pink) in both younger and older adults. In a) Bootstrap ratios are thresholded at a value of ≥3.00, which approximates a 99% confidence interval and increase from red to yellow (no regions showed reliable decrease).

### GABA+ levels and Modulation of variability

Raw GABA+/H_2_O levels (measured using MR spectroscopy) were significantly lower in older adults compared to younger adults (t (131.9) = -6.6, p = 8e-10, Cohen’s d = 1.1). Since the strongest task-condition effects on SD_BOLD_ were present in the visual cortex as were our GABA estimates, we investigated the role of Age and GABA levels in modulation of brain signal variability during visual task 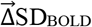 within an anatomically defined task-relevant visual mask in all subjects (see Methods). We found a single significant latent variable (p = 0.018) capturing a positive correlation between 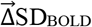 and ventrovisual GABA levels and a negative correlation between 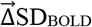 and age. As is typical when using multivariate PLS^36^, a *brainscore* was computed for each subject as the dot product between the resulting PLS model brain-pattern (i.e., a vector of brain voxel weights; see Methods) and 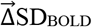 at each voxel. *Brainscores* computed within a functional mask based on the SD_BOLD_ Task-Condition model (consisting of all significant voxels from Figure 2) were highly correlated with those computed using the anatomical mask (r(133) = 0.95, p<2.2e-16)). Thus, we used the brain pattern based on the entire anatomical mask for all further analysis which allows for better reproducibility in future studies.

Both age (F(1,120) = 62.8, p < 1.3e-12, Cohen’s f = 0.71) and GABA (F(1,120) = 7.01, p = 0.009, Cohen’s f = 0.22) also remained robustly associated with visual 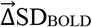 after accounting for gray-matter volume differences. Higher GABA levels were similarly associated with greater 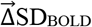 in both younger and older adults (an Age x GABA interaction was not significant, *F*(1,120) = 1.07, *p*=0.3; Figure 3a). The regions exhibiting a robust relationship (using 1000 bootstraps) between 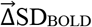 and GABA levels were in the bilateral fusiform, calcarine and lingual cortex (see Figure 3b for spatial extent and Table S2 for cluster details). These results suggest that individual differences in GABA levels play a role in how individuals modulate brain signal variability.

**Figure 3:**
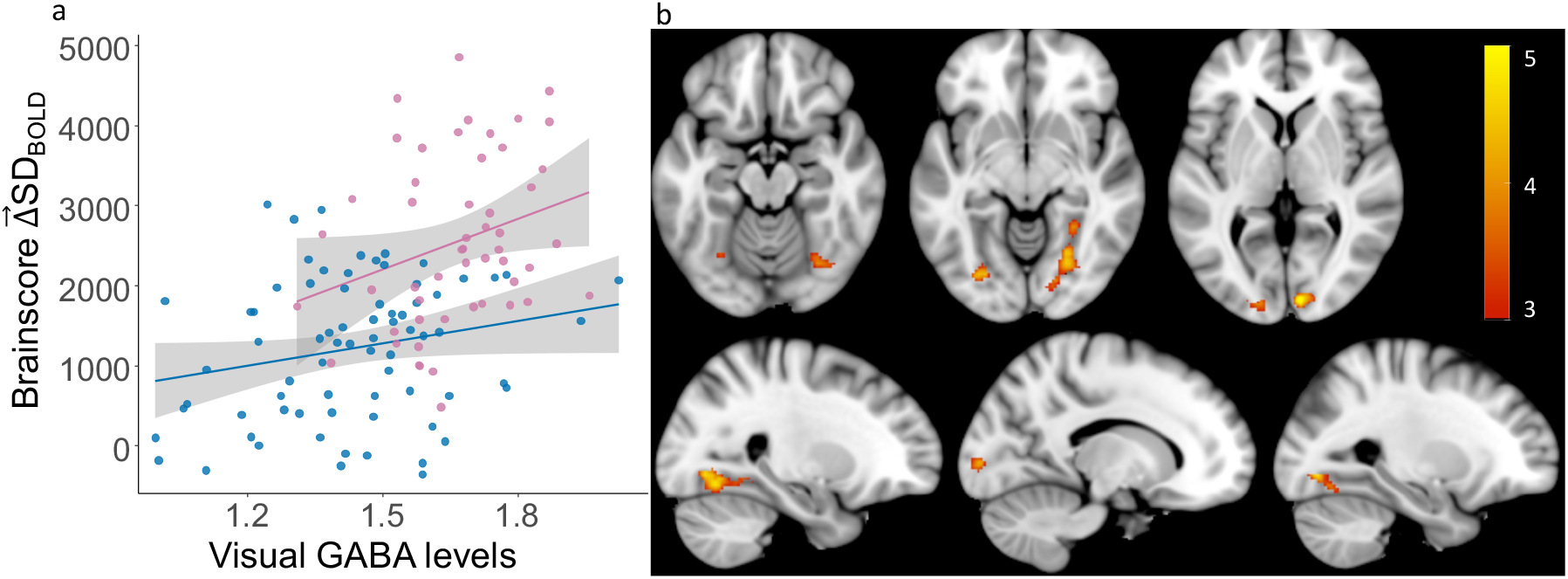
Higher GABA+ levels are associated with greater 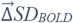. (a) Higher multivariate 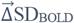 is associated with younger adult age (Cohen’s f = 0.71) and with greater GABA+ levels (Cohen’s f = 0.22) in both older (in blue) and younger adults (in pink). (b) Relationship between 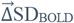 and GABA levels is robust in primary visual cortex and fusiform gyrus. Bootstrap ratios are thresholded at a value of ≥3.00, which approximates a 99% confidence interval and increase from red to yellow.

### Effect of GABA agonism on 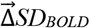

To examine a more causal role of GABA, we then administered lorazepam (a benzodiazepine known to potentiate the activity of GABA) in a subset of our participants (25 older and 20 younger adults). Masking by the same brain-pattern from the 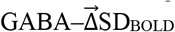 model shown in Figure 3, we estimated the influence of drug on 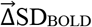 and its relationship with baseline 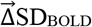 GABA levels. Consistent with a GABAergic inverted-U account, we found that baseline GABA levels measured using MRS were negatively associated with the drug-related shift in 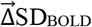 (F (1,37) = 7.85, p = 0.008, Cohen’s f = 0.42) across all subjects (see Figure 4), even after accounting for age, dosage, days between two sessions, order of sessions, and gray-matter volume differences. Age-Group did not interact with GABA level (F (1,37) = 0.9, p = 0.36). The average effect of drug was not significant on 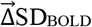 (F (1,44) = 0.04, 0.85), likely because adults with lower baseline GABA exhibited a drug-related increase in 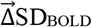, while those with higher baseline GABA showed a drug-related decrease.

**Figure 4:**
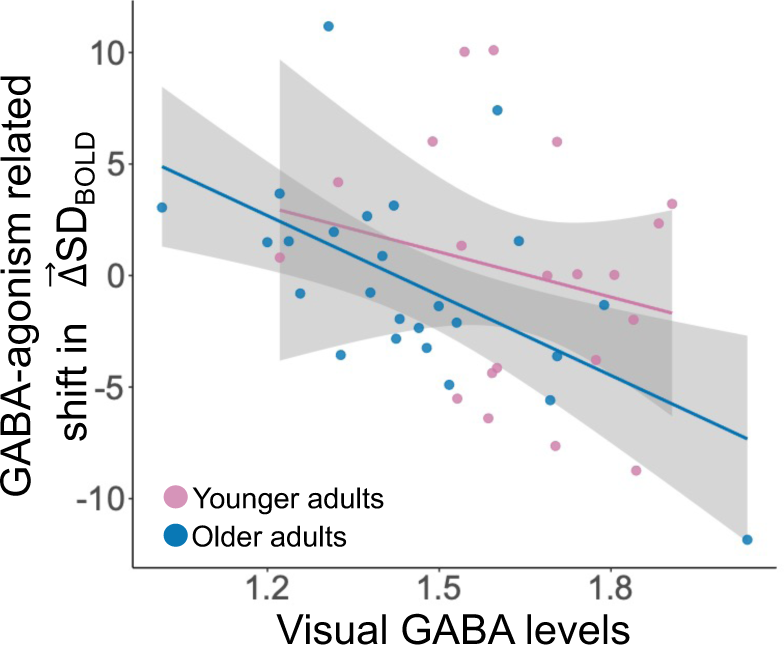
Drug-related shift in 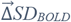 is associated with GABA+ levels. GABA agonism leads to an increase in *ΔSD*_*BOLD*_ in participants with lower GABA levels, while those with higher GABA levels exhibit a decrease in 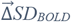 (Cohen’s f = 0.42) in both older adults (in blue) and younger adults (in pink).

### GABA+, 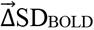, and behavior

To investigate associations between behavior and both 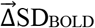 and GABA levels, a different subset of our participants (38 older and 36 younger adults) also completed four visual discrimination tasks: Buildings-in-noise, Faces-in-noise, Objects-in-noise and Scenes-in-noise. Using factor analysis, we computed a single latent factor score based on all these tasks (all loadings were positive; see Methods).

We also observed a significant Age-Group x GABA level interaction on visual discrimination behaviour, even after controlling for gray matter volume (F(1,65) = 7.13, p = 0.01, Cohen’s f = 0.3). On examining the interaction further (See Figure 5a), we found that GABA levels were more strongly associated with better visual performance in younger adults (rho(30) = 0.62, p = 0.0002, *Spearman’s correlation*) than in older adults (rho(36) = 0.37, p = 0.022). Regardless, higher GABA levels were associated with better visual performance in both younger and older adults.

**Figure 5:**
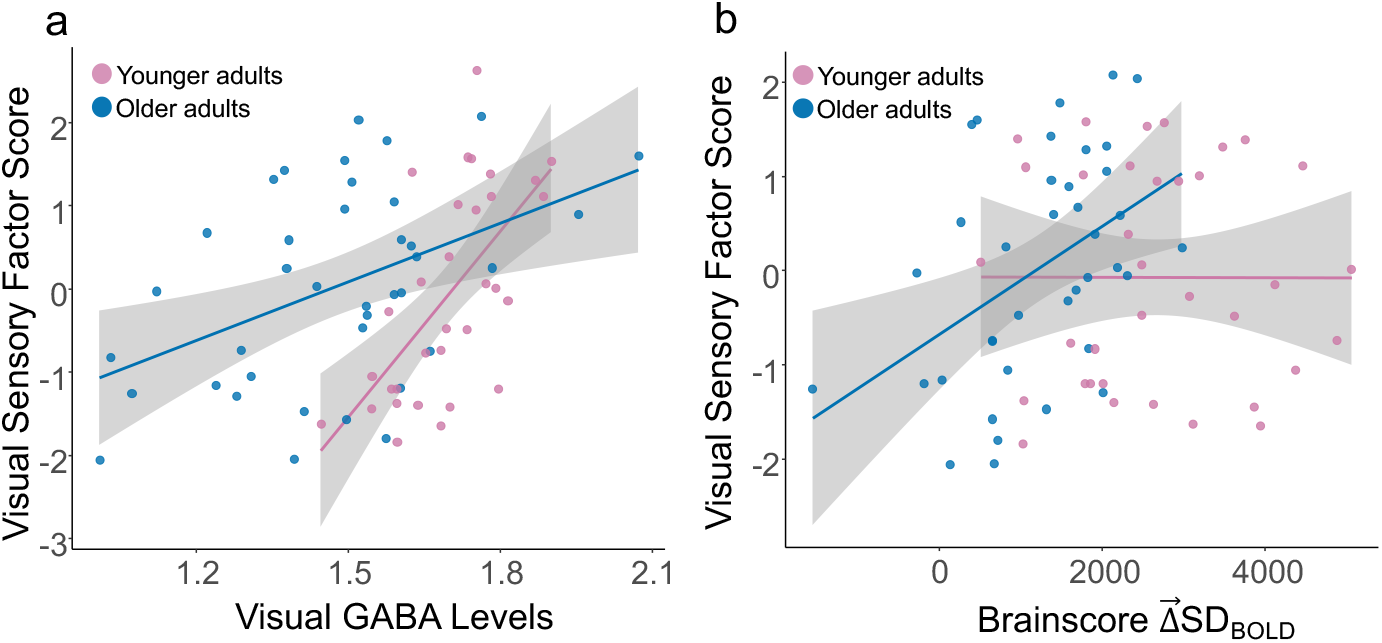
Visual sensory function, GABA+ levels, and 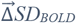. (a) Greater visual sensory scores are significantly associated with higher GABA levels in older adults (in blue) (rho(36) = 0.41) and younger adults (in pink) (rho(30) = 0.62). (b) Higher visual sensory scores were also significantly associated with 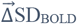 in older adults (rho(36) = 0.41) but not in younger adults.

We also found that the *Brainscores*, computed from the previously presented 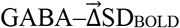 model, were significantly associated with visual performance in the older (rho(36) = 0.41, p = 0.01), but not younger group (rho(33) = 0.002, p = 0.99; see Figure 5b) and that the Age-Group x *Brainscore* interaction was significant (F(1,68) = 4.84, p = 0.03, Cohen’s f = 0.23), even after controlling for gray matter volume. Thus, both greater 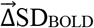 and higher GABA levels were associated with better visual discrimination performance in older adults.

Since, both 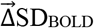 and GABA levels were associated with visual discrimination performance in older adults, we applied a hierarchical regression to compare a model explaining visual discrimination performance based on GABA levels alone (F(1,36) = 9.3, p = 0.004, Adj. R^2^ = 0.18) to one including both 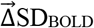(F(1, 35) = 10.42, p = 0.003) and GABA levels (F(1,35) = 7.2, p = 0.01) (Total Model: F(2,35) = 8.8, p = 0.0008, Adj. R^2^ = 0.3). We found that the model including both 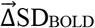 and GABA explained significant variance over and above that explained by model based on GABA alone (F(1,35) = 6.8, p = 0.01, Cohen’s f = 0.4) suggesting that 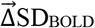 captures individual differences in visual sensory function beyond those related to differences in GABA levels in older adults.

## DISCUSSION

We report five main findings. First, brain signal variability in the visual cortex was modulated during various task conditions (greater when viewing of more complex houses vs. faces). Second, this modulation of variability 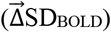 and MRS-based GABA measures in the ventrovisual cortex were both significantly lower in older adults than in younger adults. Third, individual differences in 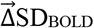 were significantly associated with GABA levels in the ventrovisual cortex. Fourth, participants with low GABA-levels tend to exhibit a drug-related increase in 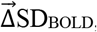, while those with high GABA-levels tended to exhibit a drug-related decrease. Fifth, both higher GABA levels and greater 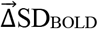 in the visual cortex were jointly associated with individual differences in visual function in older adults. We will discuss each of these findings in turn below.

### Visuo-cortical SD_BOLD_ aligns with the complexity of visual input and such alignment is lower in older adults

Consistent with prior research in older adults by Garrett et.al. (2020), we replicated the finding that BOLD signal variability (SD_BOLD_) is modulated based on the complexity of visual stimuli. SD_BOLD_ in response to more “feature-rich” house stimuli was greater than in response to simpler face stimuli in the visual cortex. These findings support the hypothesis that neural activity in the visual system should exhibit greater dynamic range in response to more complex input^37^. Indeed, greater variance in stimulus features has been argued to drive salience in the visual system^1^, which in turn could lead to an increase in resource allocation. It is hypothesized that the visual system (a) reduces resource allocation (narrowing neural dynamic range) when stimulus input is more reducible or less feature rich (here, faces), while it (b) upregulates dynamic range when stimulus input is more differentiated or feature rich (here, houses). Such upregulation of variability in response to the complexity of sensory input may reflect a well-adapted, dynamic neural network, akin to a “well-adapted organism,” as discussed by Marzen and DeDeo, (2017)^38^. Our finding that older adults were less able than younger adults to align neural variability to the complexity of visual input also converges with several previous studies showing that across-domain task-based variability modulation is increasingly muted with older adult age^4–6^. Older adults may indeed be less able to adapt to processing demands in their environment^8,39^. Beyond age differences in variability alignment, our primary goal in the present study was to better understand the mechanistic (GABAergic) basis of why this may be the case.

### The GABAergic basis of neural variability modulation

In line with our assumptions, we found that lower baseline GABA levels in the ventrovisual cortex were significantly associated with lower 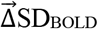 during visual processing in both young and older adults. How does the brain’s inhibitory activity (as measured by GABA levels here) play a role in modulation of brain signal variability? Computational modelling and animal research provide multiple in-roads.

Having sufficient inhibitory activity to offset excitatory activity has been found to be essential to allow artificial neural networks to operate near so-called criticality^15,40^, allowing them to visit a variety of different network states linked with higher dynamic range^16^ and brain signal variability^29^. Thus, GABA (brain’s major inhibitory neurotransmitter) plays a critical role in dictating the dynamic range of a neural network. Research in orientation selective cells in the visual cortex of cats has demonstrated that the time averaged activity of the single neuron’s spontaneous spikes in the absence of visual stimulation, is similar to the evoked pattern in presence of stimulation^41^. Thus, the dynamic range of a visual network in the absence of stimulus dictates its ability to distinctly represent visual stimuli. A flexible neural network which can sample higher number of network states should presumably represent different stimuli across the available multiple different states - leading to distinct representations of stimuli from each other characteristic of a “well-adapted organism”. Indeed, lower visual GABA levels in older adults have been associated with less differentiated response to different visual stimuli in primates^42^ and humans^17^. Our results indicate that lower GABA levels in older adults might underlie the reduced ability to align neural variability to stimulus complexity, potentially yielding impoverished stimulus differentiation in the older adult brain.

Because MR spectroscopy only captures the baseline availability of GABA, we proceeded to test the role of GABA in 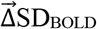 more causally by administering lorazepam, a benzodiazepine known to potentiate GABA activity at GABA_A_ receptors. Indeed, previous research in rodents noted that manipulating GABA levels in healthy rodents leads to a change in neural variability^15^. We found in humans that boosting GABA activity in participants with lower baseline GABA levels led to a boost in 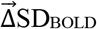 during visual processing, while those with higher baseline GABA exhibited no change or even a decrease in 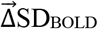 on drug. These findings provide first evidence in humans for an inverted-U account of how GABA may link to variability modulation. It is important to note that past work arguing for the presence of neurochemical inverted-U effects in humans has rarely if ever estimated baseline neurochemical levels in specific brain regions prior to pharmacological manipulation, instead typically relying on simple genotyping as a baseline estimate (e.g., COMT as a proxy for baseline dopamine^43^). We argue that MR spectroscopy may provide the easiest, regionally-specific baseline estimate of GABA for future tests of the GABAergic inverted-U.

### Higher GABA and variability modulation reflect better visual discrimination

If GABA and modulation of neural variability indeed help the brain better differentiate visual stimuli, then both should also manifest in better visual discrimination performance. Consistent with this hypothesis, we found that higher ventral visual GABA levels and higher 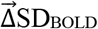 were both associated with better latent visual discrimination in-noise performance across different stimulus categories (buildings, faces, scenes, and objects). The GABA-behavior relationship was present in both age groups and is consistent with the brain maintenance in successful aging hypothesis, which suggests that older adults who maintain neurochemical levels comparable to those in younger adults experience fewer age-related impairments^44,45^.

The role of GABA in visual performance is consistent with previous research which found that GABA levels in the primary visual cortex are associated with better orientation discrimination in neurons in corresponding early visual processing areas^46^. Individual differences in GABA levels in other brain areas is also linked with better sensory performance. For example, sensorimotor GABA levels predict motor function^20,26^, auditory GABA levels predicts hearing loss^47,48^, and fronto-parietal GABA levels predict cognitive function across multiple domains assessed by higher MOCA scores^21^. We show behaviourally relevant and distinct role of GABA in the modulation of neural variability during visual processing.

Moreover, beyond previous work showing that greater task-based modulation of neural variability is associated with better behavioural performance across a variety of general cognitive domains^4,6–10^, we show that the ability to align variability to stimulus complexity 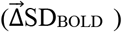 reflects visual discrimination directly, especially in older adults. Together, our results highlight the potential of targeting GABA activity in older adults with lower GABA levels to improve stimulus representations as characterized by 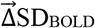, and in turn, visual processing.

We also found that within older adults, 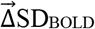 explains variance in visual task performance beyond that explained by GABA. What might underlie these individual differences in older adults? Previous research has found that not only GABA, but dopamine levels also affect the modulation of variability during visual tasks^49^. Animal research also suggests that excitatory/inhibitory (E/I) *balance* that plays a critical role in network flexibility^16^. Thus, the level of excitatory neurochemicals (e.g., glutamate) might also be an important factor in age-related changes in sensory processing. Therefore, we hypothesize that along with GABA, other age-related neurochemical changes might also underly individual differences in 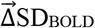 and visual processing. A direct comparison of the role of dopamine, GABA, and glutamate would be ideal in future work.

The current research focused on understanding the role of GABA and 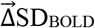 in visual processing. However, previous research has highlighted the role of higher 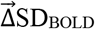 in better performance in other cognitive domains such as processing speed^6^ and memory^49,50^, while lower SD_BOLD_ has been related to age-linked pathologies like Alzheimer’s disease^51,52^. Separately, researchers have also found a link between dysfunction in GABAergic signaling and AD^53^ as well as poor cognitive functions^21^ as described above. Investigating if age-related changes in GABA and 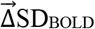 underlie these pathologies and cognitive decline might hold promise for developing new pharmacological targets to alleviate age-linked impairments.

## Conclusion

Our results suggest that GABA plays a key role in how humans of different ages modulate neural variability to adapt to the complexity of the visual world. Interventions targeting inhibitory systems could serve to slow both sensory and variability-based declines characteristic of older adults, a key goal for future work.

## MATERIALS and METHODS

This data was collected as part of the Michigan Neural Distinctiveness (MiND) study. Here we only describe the portions of the study that are relevant to this analysis. For details about the entire study protocol see (Gagnon et al., 2019^54^). The ethical approval for the study was granted by the Institutional Review Board of The University of Michigan (HUM00103117).

### Participants

We analyzed the data from 58 young (age 18-29 years) and 77 older (age 65 and above) adults who completed the entire MiND study before the start of the COVID-19 pandemic. All participants were recruited from Ann Arbor and the surrounding area, were right-handed, native English speakers, and had normal or corrected to normal vision. We screened out participants who scored 23 or lower on the Montreal Cognitive Assessment (MOCA)^55^. All the sessions described below took place at the University of Michigan’s Functional MRI Laboratory, Ann Arbor, Michigan.

### Power Analysis

Garrett et.al. (2020) found a correlation of r=0.47 between modulation of visual variability and behavioral performance. In our convenience sample of those who completed the study before the pandemic and who completed the behavioral tasks, we had 80% power to detect a correlation of r=0.47 in each age-group alone.

### Session Design

After completing an initial telephone screening interview and being determined eligible, all subjects participated in three sessions, each on a separate day. Session 1 only involved cognitive and behavioral testing, session 2 included behavioral testing and a functional magnetic resonance imaging (fMRI) scan, and session 3 only involved a Magnetic Resonance Spectroscopy (MRS) scan (See Figure 1 for summary).

### Behavioral Testing

A subset of participants (38 older and 36 younger adults) completed four visual tasks in noise, administered on a Dell laptop with a 15.6-inch screen using the Psychophysics Toolbox. All tasks consisted of trials in which a 500 ms fixation cross was followed by a black and white picture in dynamic Gaussian noise for 500 ms followed by a response screen. The next trial began after the response. The order of the stimulus presentation was pseudorandomized but was the same across participants. Each task began with 4 practice trials with feedback and was followed by 50 scored trials without feedback. The tasks followed a staircase procedure – when a participant made three correct responses in a row, the level of noise was increased. Following an incorrect response, the level of noise was decreased. There were a total of 15 levels of Gaussian noise, and each task started at the 5^th^ level of noise. The dependent measure was the average level of noise presented for the last 40 trials. Thus, a higher score represents better performance.

#### a. Buildings in Noise (BIN)

The stimulus picture was either a house (50% of trials) or an apartment (50% of trials). Participants were asked to press “1” with their left index finger if they thought the picture was a house and “0” with their right index finger if they thought the picture was an apartment. Stimuli were from Park et al., (2004)^56^.

#### b. Faces in Noise (FIN)

The stimulus picture was either a male (50% of trials) or female (50% of trials) face. Participants pressed “1” if they thought the picture was a male face and “0” if they thought the picture was a female face. Stimuli were from Gold et al., (1999)^57^.

#### c. Objects in Noise (OIN)

The stimulus picture was either an office item, such as a stapler (50% of trials), or a food item, such as a hamburger (50% of trials). Participants pressed “1” for an office item and “0” for a food item. Object stimuli were taken from Brady et al., (2008)^58^.

#### d. Scenes in Noise (ScIN)

The stimulus picture was either an urban (50% of trials) or nature (50% of trial) scene. Participants pressed “1” if they thought the picture was an urban scene and “0” for a nature scene. Scene images were from Zhou et al., (2018)^59^.

Using the MATLAB function *factoran* we computed a single visual sensory factor based on these four tasks. The loadings of the tasks on the factor were 0.3 (buildings), 0.8 (faces), 0.5 (objects) and 0.2 (scenes). Higher factor scores reflect better performance. Using principal component analysis (PCA) only one component had an eigen value > 1, explaining 40% of the variance, and the corresponding weights were 0.53 (buildings), 0.8 (faces), 0.7 (objects) and 0.42 (scenes). The scores on this first component were correlated at r(72) = 0.88, we thus present results based on factor score.

### fMRI Session

Functional MRI data was collected using a 3T General Electric Discovery Magnetic Resonance System with a volumetric quadrature bird cage head coil and two 32- channel receive arrays. The functional scan parameters were as follows: T2*-weighted images using a 2D Gradient Echo pulse sequence; Repetition Time (TR) = 2000 ms; Echo Time (TE) = 30 ms; flip angle = 90°; Field of View (FOV) = 220 × 220 mm; 185 volumes; 43 axial slices; thickness = 3 mm, no spacing; and collected in an interleaved bottom-up sequence. The total acquisition time for the visual task scan was 6 minutes 10 seconds.

The task consisted of six 20-second blocks of images of male faces and six 20-second blocks of images of houses, presented in a pseudorandomized order and interleaved with twelve 10-second blocks of a fixation cross. Each block consisted of the stimulus presented for 500 ms with an interstimulus interval (ISI) of 500 ms. The block order was the same for all the participants. This was a passive viewing task but to ensure participant attention, we presented rare target trials once every minute (6 in total). The target trial were images of female faces for the face blocks, and images of apartment buildings for the house blocks. Participants were instructed to press a button with their right index finger every time they saw a target trial.

### MRS Session

Magnetic resonance spectroscopy (MRS) scanning was completed on a different day using the same MRI scanner described above. This session lasted approximately 1.5 hours, during which we collected another T1-weighted structural image and MRS data. The T1-weighted image was obtained using the same parameters and sequence used during the fMRI session. The MRS data was obtained from 3cm x 3cm x 3cm voxels placed in the left and right ventrovisual cortex. The voxel placement was guided by the person-specific task-based fMRI activations, such that the voxels were centered roughly at the peak of activation for a face and house viewing vs. fixation contrast, separately for each participant. We used a MEGA-PRESS sequence with the following parameters to obtain MR spectra: TE=68ms (TE1=15ms, TE2=53ms), TR=1.8sec, spec. width=2kHz, Frequency selective editing pulses (14ms) applied at 1.9ppm (ON) & 7.46 ppm (OFF).

### Drug session

A subset of participants (20 young and 25 older adults) underwent two functional MRI sessions instead of one, one after taking a low dose benzodiazepine (lorazepam) and one after taking a placebo pill. The functional scanning parameters were identical to those described above. The pills were given approximately 1 hour before the session and the order of the sessions (on and off drug) was counterbalanced across participants. During the drug session, participants were administered a 0.5 or 1 mg oral dose of lorazepam (a benzodiazepine). The dosage was assigned randomly. Participants were not told which pill they received on which day (they were blind to the drug administration order).

### fMRI Data Preprocessing

The fMRI data were preprocessed and analyzed using a combination of FMRIB Software Library (FSL), SPM12 and MATLAB-based scripts. The first 5 volumes of each scan were discarded. Heart rate was collected via a pulse oximeter placed on the left middle finger and the data was physio corrected during preprocessing. We performed 1^st^-level preprocessing using FSL-FEAT^60^ with default parameters for motion correction, normalization, and smoothing (7mm). We used the SPM12 function spm_detrend to remove linear, quadratic and cubic trends in the time series and also applied a Butterworth filter (0.01-0.1Hz). We then ran FSL MELODIC to perform Independent Component Analysis (ICA). Three separate raters identified noise components through manual visual inspection^61^. These components reflected noise related to sinus activity, vascular and ventricle activations, and motion. We then removed the components identified as noise by at least two of the three raters using the FSL regfilt function. Subsequently, we performed linear registration of the functional and anatomical images of each participant and the MNI152 template using the FSL FLIRT function.

### Quantification of Brain signal variability (SD_BOLD_)

After preprocessing the fMRI data, the standard deviation in the fMRI signal during each of the conditions (faces and houses) was computed at each voxel for each participant as was done in previous work^3^. Modulation of variability 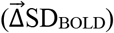 was computed as the difference between the two conditions (SD_BOLD-HOUSES_ – SD_BOLD- FACES_) at each of the voxels in a grey matter volume mask. Similarly, 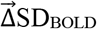 on drug was computed for the subset of participants who were administered lorazepam.

### Quantification of GABA levels

We used Gannet 3.0^62^, a MATLAB based toolbox, to estimate GABA+/Water levels based on the MEGA- PRESS difference spectra in each of the MRS voxels. All the time-domain data were phase corrected and frequency corrected using spectral registration and filtered with 3- Hz exponential line broadening and zero-filled by a factor of 16. The GABA levels were scaled to water and expressed in institutional units by Gannet. Gannet quantifies GABA levels by fitting a five-parameter Gaussian model to the MR spectrum between 2.19 and 3.55 ppm while the water peak is modelled using a Gaussian-Lorentzian function. The MEGA-PRESS editing scheme also results in excitation of coedited macromolecules (MM), which can contribute up to 45% to the edited signal around 3ppm overlapping with the GABA peak. Thus, all GABA values are reported as GABA+ (i.e., GABA + MM) in the present study.

Gannet’s integrated voxel-to-image co-registration procedure produces a binary mask of the MRS voxel. Using an SPM-based segmentation function, Gannet estimates the tissue composition (voxel fractions containing Cerebrospinal Fluid (CSF), Grey matter (GM) and white matter (WM)). Gannet then estimates a tissue- corrected GABA+ value that accounts for the fraction of grey matter, white matter, and CSF in each MRS voxel as well as the differential relaxation constants and water visibility in the different tissue types^63^. Based on a quality control check of the spectra, we flagged and discarded three right and seven left GABA values. GABA measures between right and left ventrovisual voxel were correlated at r(134) = 0.57 (p<0.0001). Thus, we computed an average GABA+ measure for each participant. One young participant had both right and left ventrovisual GABA values flagged and was excluded from the analysis.

### Statistical Analysis

To estimate the effects of task-condition on SD_BOLD_ and the effect of age and GABA on 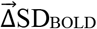 we employed multivariate partial least squares (PLS) analyses^64^.

1. To estimate the effects of task-condition on SD_BOLD_ we used a task-PLS. In this method, first a between- subject covariance (COV) matrix is computed between house and face conditions and each voxel’s SD_BOLD_. Then a left singular vector of experimental condition weights (U) is estimated, along with a right singular vector of brain voxel weights (V) and a diagonal matrix of singular values (S). Significance of the detected relations is assessed using 1000 permutation tests of the singular value corresponding to the latent variable (LV). This resulted in two LVs of which only one was significant and represented greater variability during the house condition than during the face condition. *Brainscore* was computed separately for each condition as the dot product of brain voxel weights and each subjects’ SD_BOLD-HOUSES_ and SD_BOLD-FACES_.
2. We employed rank-based Behavior-PLS for investigating effects of age and GABA levels on 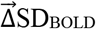 in a visual anatomical mask within our sample of 134 subjects (after excluding the one subject whose GABA estimates were flagged). The visual anatomical mask is a broad mask that contained the occipital pole, lingual gyrus, inferior division of lateral occipital cortex, the temporooccipital part of the inferior and middle temporal gyrus, temporal occipital fusiform cortex, occipital fusiform gyrus, and parahippocampal gyrus from the Harvard-Oxford atlas in FSL. A between- subject correlation matrix (CORR) was computed between each voxel’s 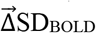 (i.e. SD_BOLD-HOUSES_ – SD_BOLD-FACES_) within this mask and both – 1) MRS- based average ventrovisual GABA measures and 2) self-reported age (in years). Then, CORR was decomposed using singular value decomposition (SVD). This decomposition produced a matrix of behavior weights (U), a matrix of brain voxel weights (V), and a diagonal matrix of singular values (S). A single significant latent variable captured the activity pattern depicting the brain regions that show the strongest relationship of 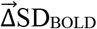 with both age and GABA. The behavior weights (U) of this LV suggest lower GABA levels and greater age were associated with lower 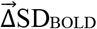. We obtained a summary measure of each participant’s expression of this LV’s spatial pattern (a within-person “*Brainscore*”) by taking a dot product of the brain weights (V) with 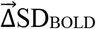 on placebo (and on drug for the subset that received the manipulation). This *Brainscore* was also used for investigating the role of 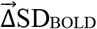 on visual discrimination task.

In both models, we used a bootstrapping procedure (1000 bootstrapped resamples) to reveal the robustness of voxel weights. By dividing each voxel’s weight by its bootstrapped standard error, we obtained “bootstrap ratios” (BSRs) as estimates of robustness. We thresholded BSRs at values of ±3.00 (∼99.9% confidence interval). Statistical analyses were conducted using R^65^. Figures were plotted using the ggplot2 package^66^ and the lme4 package^67^ was used to perform the linear mixed effects analysis. We used a Cook’s Distance greater than 4/sample size to flag and remove outliers from each of mixed effects models reported in the results. See supplementary for full sample results.

## REFERENCES

1. Hermundstad, A. M. et al. Variance predicts salience in central sensory processing. eLife 3, e03722 (2014).

2. Orbán, G., Berkes, P., Fiser, J. & Lengyel, M. Neural Variability and Sampling-Based Probabilistic Representations in the Visual Cortex. Neuron 92, 530–543 (2016).

3. Garrett, D. D., Epp, S. M., Kleemeyer, M., Lindenberger, U. & Polk, T. A. Higher performers upregulate brain signal variability in response to more feature-rich visual input. NeuroImage 217, 116836 (2020).

4. Garrett, D. D., Kovacevic, N., McIntosh, A. R. & Grady, C. L. Blood Oxygen Level-Dependent Signal Variability Is More than Just Noise. J Neurosci 30, 4914–4921 (2010).

5. Garrett, D. D., Kovacevic, N., McIntosh, A. R. & Grady, C. L. The Importance of Being Variable. J Neurosci 31, 4496–4503 (2011).

6. Garrett, D. D., Kovacevic, N., McIntosh, A. R. & Grady, C. L. The Modulation of BOLD Variability between Cognitive States Varies by Age and Processing Speed. Cereb Cortex 23, 684–693 (2013).

7. Grady, C. L. & Garrett, D. D. Brain signal variability is modulated as a function of internal and external demand in younger and older adults. Neuroimage 169, 510–523 (2018).

8. Waschke, L., Kloosterman, N. A., Obleser, J. & Garrett, D. D. Behavior needs neural variability. Neuron 109, 751–766 (2021).

9. Rieck, J. R., Rodrigue, K. M., Boylan, M. A. & Kennedy, K. M. Age-related reduction of BOLD modulation to cognitive difficulty predicts poorer task accuracy and poorer fluid reasoning ability. NeuroImage 147, 262–271 (2017).

10. Rieck, J. R., DeSouza, B., Baracchini, G. & Grady, C. L. Reduced modulation of BOLD variability as a function of cognitive load in healthy aging. Neurobiology of Aging 112, 215–230 (2022).

11. Jones, E. G. GABAergic Neurons and Their Role in Cortical Plasticity in Primates. Cereb Cortex 3, 361– 372 (1993).

12. Hensch, T. K. et al. Local GABA circuit control of experience-dependent plasticity in developing visual cortex. Science 282, 1504–1508 (1998).

13. Fagiolini, M. et al. Specific GABAA circuits for visual cortical plasticity. Science 303, 1681–1683 (2004).

14. Monteforte, M. & Wolf, F. Dynamical entropy production in spiking neuron networks in the balanced state. Physical review letters 105, 268104 (2010).

15. Shew, W. L., Yang, H., Yu, S., Roy, R. & Plenz, D. Information Capacity and Transmission Are Maximized in Balanced Cortical Networks with Neuronal Avalanches. J. Neurosci. 31, 55–63 (2011).

16. Agrawal, V. et al. Robust entropy requires strong and balanced excitatory and inhibitory synapses. Chaos 28, 103115 (2018).

17. Chamberlain, J. D. et al. GABA levels in ventral visual cortex decline with age and are associated with neural distinctiveness. Neurobiology of Aging 102, 170–177 (2021).

18. Lalwani, P. et al. Neural distinctiveness declines with age in auditory cortex and is associated with auditory GABA levels. NeuroImage 201, 116033 (2019).

19. Simmonite, M. et al. Age-Related Declines in Occipital GABA are Associated with Reduced Fluid Processing Ability. Acad Radiol (2018) doi:10.1016/j.acra.2018.07.024.

20. Cassady, K. et al. Sensorimotor network segregation declines with age and is linked to GABA and to sensorimotor performance. NeuroImage 186, 234– 244 (2019).

21. Porges, E. C. et al. Frontal Gamma-Aminobutyric Acid Concentrations Are Associated With Cognitive Performance in Older Adults. Biol Psychiatry Cogn Neurosci Neuroimaging 2, 38–44 (2017).

22. Chalavi, S. et al. The neurochemical basis of the contextual interference effect. Neurobiol. Aging 66, 85–96 (2018).

23. Hermans, L. et al. GABA levels and measures of intracortical and interhemispheric excitability in healthy young and older adults: an MRS-TMS study. Neurobiology of Aging 65, 168–177 (2018).

24. Gao, F. et al. Edited magnetic resonance spectroscopy detects an age-related decline in brain GABA levels. Neuroimage 78, 75–82 (2013).

25. Cuypers, K., Maes, C. & Swinnen, S. P. Aging and GABA. Aging (Albany NY) 10, 1186–1187 (2018).

26. Levin, O. et al. Sensorimotor cortex neurometabolite levels as correlate of motor performance in normal aging: evidence from a 1H-MRS study. NeuroImage 202, 116050 (2019).

27. Porges, E. C., Jensen, G., Foster, B., Edden, R. A. & Puts, N. A. The trajectory of cortical GABA across the lifespan, an individual participant data meta-analysis of edited MRS studies. eLife 10, e62575 (2021).

28. Porges, E. C. et al. Impact of tissue correction strategy on GABA-edited MRS findings. NeuroImage 162, 249–256 (2017).

29. Lalwani, P., Garrett, D. D. & Polk, T. A. Dynamic Recovery: GABA Agonism Restores Neural Variability in Older, Poorer Performing Adults. J. Neurosci. 41, 9350–9360 (2021).

30. Młynarski, W. F. & Hermundstad, A. M. Adaptive coding for dynamic sensory inference. eLife 7, e32055 (2018).

31. Northoff, G. & Tumati, S. “Average is good, extremes are bad” – Non-linear inverted U-shaped relationship between neural mechanisms and functionality of mental features. Neuroscience & Biobehavioral Reviews 104, 11–25 (2019).

32. Riesenhuber, M. & Poggio, T. Hierarchical models of object recognition in cortex. Nature neuroscience 2, 1019–1025 (1999).

33. Serre, T., Oliva, A. & Poggio, T. A feedforward architecture accounts for rapid categorization. Proceedings of the national academy of sciences 104, 6424–6429 (2007).

34. Serre, T., Wolf, L., Bileschi, S., Riesenhuber, M. & Poggio, T. Robust object recognition with cortex-like mechanisms. IEEE transactions on pattern analysis and machine intelligence 29, 411–426 (2007).

35. Serre, T. et al. A theory of object recognition: computations and circuits in the feedforward path of the ventral stream in primate visual cortex. (2005).

36. Krishnan, A., Williams, L. J., McIntosh, A. R. & Abdi, H. Partial Least Squares (PLS) methods for neuroimaging: a tutorial and review. Neuroimage 56, 455–475 (2011).

37. Van Hateren & h, J. Real and optimal neural images in early vision. Nature 360, 68–70 (1992).

38. Marzen, S. E. & DeDeo, S. The evolution of lossy compression. Journal of The Royal Society Interface 14, 20170166 (2017).

39. Garrett, D. D. et al. Moment-to-moment brain signal variability: A next frontier in human brain mapping? Neurosci Biobehav Rev 37, 610–624 (2013).

40. Poil, S.-S., Hardstone, R., Mansvelder, H. D. & Linkenkaer-Hansen, K. Critical-State Dynamics of Avalanches and Oscillations Jointly Emerge from Balanced Excitation/Inhibition in Neuronal Networks. J. Neurosci. 32, 9817–9823 (2012).

41. Buzsaki, G. Rhythms of the Brain. (Oxford University Press, 2006).

42. Leventhal, A. G., Wang, Y., Pu, M., Zhou, Y. & Ma, Y. GABA and its agonists improved visual cortical function in senescent monkeys. Science 300, 812– 815 (2003).

43. Bäckman, L., Nyberg, L., Lindenberger, U., Li, S.-C. & Farde, L. The correlative triad among aging, dopamine, and cognition: Current status and future prospects. Neuroscience & Biobehavioral Reviews 30, 791–807 (2006).

44. Nyberg, L., Lövdén, M., Riklund, K., Lindenberger, U. & Bäckman, L. Memory aging and brain maintenance. Trends in Cognitive Sciences 16, 292– 305 (2012).

45. Bruffaerts, R., Tyler, L. K., Shafto, M., Tsvetanov, K. A. & Clarke, A. Perceptual and conceptual processing of visual objects across the adult lifespan. Sci Rep 9, 13771 (2019).

46. Kurcyus, K. et al. Opposite Dynamics of GABA and Glutamate Levels in the Occipital Cortex during Visual Processing. J. Neurosci. 38, 9967–9976 (2018).

47. Gao, F. et al. Decreased auditory GABA+ concentrations in presbycusis demonstrated by edited magnetic resonance spectroscopy. Neuroimage 106, 311–316 (2015).

48. Dobri, S. G. J. & Ross, B. Total GABA level in human auditory cortex is associated with speech-in-noise understanding in older age. NeuroImage 225, 117474 (2021).

49. Garrett, D. D. et al. Dynamic regulation of neural variability during working memory reflects dopamine, functional integration, and decision-making. 2022.05.05.490687 Preprint at https://doi.org/10.1101/2022.05.05.490687 (2022).

50. Garrett, D. D. et al. Amphetamine modulates brain signal variability and working memory in younger and older adults. PNAS 112, 7593–7598 (2015).

51. Millar, P. R. et al. Evaluating resting-state BOLD variability in relation to biomarkers of preclinical Alzheimer’s disease. Neurobiology of Aging 96, 233–245 (2020).

52. Zhang, L. et al. Distinct BOLD variability changes in the default mode and salience networks in Alzheimer’s disease spectrum and associations with cognitive decline. Sci Rep 10, 6457 (2020).

53. Guzmán, B. C.-F. et al. The GABAergic system as a therapeutic target for Alzheimer’s disease. Journal of Neurochemistry 146, 649–669 (2018).

54. Gagnon, H. et al. Michigan Neural Distinctiveness (MiND) study protocol: investigating the scope, causes, and consequences of age-related neural dedifferentiation. BMC Neurol 19, 61 (2019).

55. Carson, N., Leach, L. & Murphy, K. J. A re-examination of Montreal Cognitive Assessment (MoCA) cutoff scores. Int J Geriatr Psychiatry 33, 379–388 (2018).

56. Park, D. C. et al. Aging reduces neural specialization in ventral visual cortex. Proc. Natl. Acad. Sci. U.S.A. 101, 13091–13095 (2004).

57. Gold, J., Bennett, P. J. & Sekuler, A. B. Signal but not noise changes with perceptual learning. Nature 402, 176–178 (1999).

58. Brady, T. F., Konkle, T., Alvarez, G. A. & Oliva, A. Visual long-term memory has a massive storage capacity for object details. PNAS 105, 14325–14329 (2008).

59. Zhou, B., Lapedriza, A., Khosla, A., Oliva, A. & Torralba, A. Places: A 10 Million Image Database for Scene Recognition. IEEE Transactions on Pattern Analysis and Machine Intelligence 40, 1452–1464 (2018).

60. Woolrich, M. W., Ripley, B. D., Brady, M. & Smith, S. M. Temporal Autocorrelation in Univariate Linear Modeling of FMRI Data. NeuroImage 14, 1370–1386 (2001).

61. Kelly, R. E. et al. Visual inspection of independent components: defining a procedure for artifact removal from fMRI data. J. Neurosci. Methods 189, 233–245 (2010).

62. Edden, R. A. E., Puts, N. A. J., Harris, A. D., Barker, P. B. & Evans, C. J. Gannet: A batch-processing tool for the quantitative analysis of gamma-aminobutyric acid–edited MR spectroscopy spectra. J Magn Reson Imaging 40, 1445–1452 (2014).

63. Harris, A. D., Puts, N. A. J. & Edden, R. A. E. Tissue correction for GABA-edited MRS: Considerations of voxel composition, tissue segmentation, and tissue relaxations. J Magn Reson Imaging 42, 1431– 1440 (2015).

64. McIntosh, A. R., Bookstein, F. L., Haxby, J. V. & Grady, C. L. Spatial pattern analysis of functional brain images using partial least squares. Neuroimage 3, 143–157 (1996).

65. Team, R. C. R: A language and environment for statistical computing. (2013).

66. Wickham, H. ggplot2: Elegant Graphics for Data Analysis. (Springer, 2016).

67. Bates, D., Sarkar, D., Bates, M. D. & Matrix, L. The lme4 package. R package version 2, 74 (2007).

