## Supplementary Materials for "Modulation of neural variability: Age-related reduction, GABAergic basis, and behavioral implications"

### Modulation of neural variability: Age-related reduction, neurochemical cause, and behavioral consequences

#### SUPPLEMENTARY MATERIALS

##### Supplementary Results

*HMAX Analysis of Visual Stimuli*

Physiological studies in non-human primates over recent decades have demonstrated that the receptive fields of neurons increase in both size and complexity as we move anteriorly along the ventral visual pathway. These insights are reflected in the biologically inspired, openly available HMAX feedforward model of visual recognition (code source: http://maxlab.neuro.georgetown.edu/hmax.html). The two earliest layers in this model (S1 and C1) correspond to neurons in primary visual cortex (V1) and the next two layers (S2 and C2) correspond to neurons in extrastriate visual areas (V2/V4). Following Garrett et al. (2020), we used this model to objectively estimate the visual complexity of the two stimulus categories presented during our visual task (houses vs. faces). This model can also generate predictions about which cortical regions should be most sensitive to differences in stimulus complexity in our specific stimulus set. We focused our analyses on the C1 and C2 layers that aggregate the responses of cells in the S1 and S2 layers. To anticipate the results, we found that houses were more "feature-rich" than faces in both C1 (corresponding to V1) and C2 layers (corresponding to extrastriate regions), as described in detail below.

Layers in the first layer (S1) are modelled using Gabor functions with 16 different size filters (ranging from 7 to 37) corresponding to n x n pixel neighborhoods for four different orientations (Default: -45°, 0°, 45°, 90°). For each orientation and filter size, a model was fit to each image in overlapping windows (50% overlap) resulting in a simple cell response map for all positions within the input image. At the next layer (complex cells in C1), the maximum activity over S1 units corresponding to each orientation is computed separately. Since 16 filter sizes were used, taking the maximum over neighboring pairs of filters results in eight "scale bands". The scale band index corresponds to the spatial neighborhood of S1 cells over which outputs are pooled. For each of the 8 scale-bands and 4 orientations, we calculated a median within-image C1 activation value for each image and then standardized them by computing z-scores. Using t-tests, we compared these within-image median values across the two stimulus categories. The results of these 8 (scales) × 4 (orientations) independent sample t-tests (Figure S1A and S1C) indicate that houses consistently produced a larger median C1 activation value than faces across all receptive field sizes.

In the third layer (S2), a template-matching approach is used. The receptive fields of the S2 cells correspond to a set of universal prototype templates derived from a library of naturalistic stimuli and their activation is computed based on the Euclidean distance between incoming C1 activity from all four orientations and the stored prototype for that S2 cell. For each prototype, an S2 map is computed across all positions at each of the 8 scale bands. The final layer (C2) then takes a global maximum over all scales and positions for each S2 map separately for different neighborhood (patch) sizes. We computed the median activation for each C2 neighborhood size separately and then standardized the results using z-scores. We then compared median activation by faces and houses using eight independent sample t-tests (one for each patch size). We found that house stimuli showed greater median activity compared with faces across different patch sizes (Figure S1B and S1D).

*GABA estimates with tissue-corrections*

GABA+ levels were also significantly lower in older adults compared to younger adults after correcting for alpha tissue-composition differences ($t(131.9) = -3.13$, $p = 0.002$, Cohen's $d = 0.53$) (Porges et.al., 2017). These results suggest that the differences in tissue composition only partially explain the observed lower GABA levels in older adults. Raw GABA+/$H_2O$ levels were highly correlated with alpha tissue composition-corrected GABA levels ($r(131) = 0.88$, $p < 2.2e\text{-}16$). All results presented in the manuscript were thus based on the raw GABA+/$H_2O$ levels.

*Results without excluding outliers*

No outliers were determined using Cook's Distance (=4/sample size) in model examining role of task on $SD_{BOLD}$ or role of baseline GABA in drug-related shift in $\vec{\Delta}SD_{BOLD}$. Linear model investigating role of

Age and GABA on $\vec{\Delta}SD_{BOLD}$ found similar effects when computed without excluding outliers (presented in main results). Both age ($F(1,129) = 54.73$, $p < 1.6e\text{-}11$, Cohen's $f = 0.62$) and GABA levels ($F(1,129) = 7.03$, $p = 0.009$, Cohen's $f = 0.21$) had a significant effect on this *Brainscore* even after accounting for gray-matter volume differences. Higher GABA levels were associated with greater $\Delta SD_{BOLD}$ in both older and younger adults and the Age x GABA interaction was not significant ($F(1,129) = 0.0004$, $p=0.98$).

The model investigating role of GABA in visual discrimination found that GABA had a significant effect on visual latent score after controlling for age and gray matter volume ($F(1,69) = 9.7$, $p = 0.003$, Cohen's $f = 0.34$). However, the Age-Group x GABA interaction ($F(1,69) = 0.74$, $p = 0.39$) was not significant. There were 4 young outliers with Cook's Distance greater than 0.054 (4/sample size). Nonetheless, the relationship between GABA and visual performance was significant in both age-groups, with and without outliers.

The *Brainscores* computed from GABA–$\vec{\Delta}SD_{BOLD}$ model were significantly associated with the latent visual score without excluding the outlier ($F(1,69) = 4.2$, $p = 0.04$, Cohen's $f = 0.21$). The Age-Group x *Brainscore* interaction however showed a trend towards significance ($F(1,69) = 3.6$, $p = 0.06$). One young outlier subject was determined as an outlier using Cook's Distance greater than 0.054 (4/sample size) and presented in the main results. Nonetheless, the correlation was significant only in older adults after controlling for age and gray matter volume ($rho(37) = 0.41$, $p = 0.01$) but not in younger adults with ($rho(34) = -0.01$, $p = 0.95$) and without the outlier ($r(33) = -0.003$, $p = 0.99$).

#### Supplementary Figure

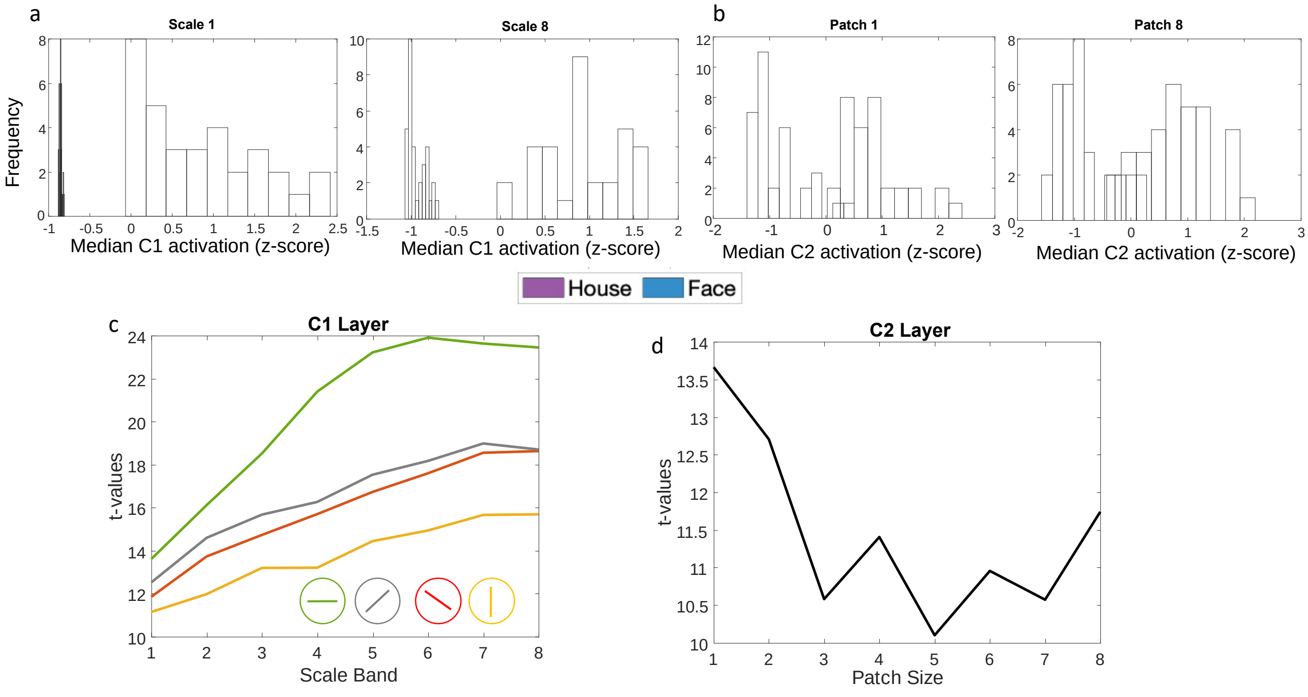


*Fig S1. Example C1 and C2 activation distributions to house and face stimuli. A) Z-scored median activation at C1 for all images in the two stimulus categories (faces in blue, houses in purple) for one orientation and two different scales. B) Z-scored median activation at C2 for all images in both stimulus categories for two different patch sizes. C) t-values comparing house vs. face median activation in layer C1 across four different orientations and 8 scale bands (from smallest to largest receptive field). D) t-values comparing house vs. face median activation in layer C2 for 8 different patch sizes. All p-values for the t-tests are less than 0.001.*

**Supplementary Table**

**Table S1**
Brain regions that exhibited a reliable association between task-condition and $SD_{BOLD}$.

| Cluster Number | MNI Co-ordinates | | | Peak Threshold (BSR) | Cluster Size (in 2mm voxels) | Cortical Region Label based on Harvard Oxford Cortical Atlas |
|---|---|---|---|---|---|---|
| | X | Y | Z | | | |
| 1 | 26 | -44 | -18 | 16.71 | 18565 | Temporal Occipital Fusiform Cortex (Includes parts of Occipital Pole (AAL Label: Calcarine, Lingual), Lingual Gyrus, Temporal Fusiform Cortex, Occipital Fusiform Gyrus, posterior Parahippocampal Gyrus etc. |

**Table S2**
Brain regions that exhibited a reliable association between age, GABA and $\vec{\Delta}SD_{BOLD}$.

| Cluster Number | MNI Co-ordinates | | | Peak Threshold (BSR) | Cluster Size (in 2mm voxels) | Cortical Region Label based on Harvard Oxford Cortical Atlas |
|---|---|---|---|---|---|---|
| | X | Y | Z | | | |
| 1 | -28 | -78 | -6 | 5.35 | 118 | (L) Occipital Fusiform Gyrus |
| 2 | -10 | -92 | 2 | 5.05 | 124 | Bilateral Occipital Pole (AAL Label: Calcarine and Lingual) |
| 3 | 24 | -70 | -8 | 4.69 | 199 | (R) Occipital Fusiform Gyrus |
